# Myelin Basic Protein Binding Is Modulated by Leaflet Asymmetry and Lipid Composition

**DOI:** 10.64898/2025.12.19.695364

**Authors:** Julio M. Pusterla, Alexandros Koutsioubas, Nicolò Paracini, Philipp Gutfreund, Maximilian W. A. Skoda, Stephen Hall, Igor Graf von Westarp, Irina Apanasenko, Marina Cagnes, Stephan Förster, Andreas M. Stadler

**Author notes:** **Corresponding Authors:** Julio M. Pusterla, Andreas M. Stadler. European Spallation Source ERIC, P.O. Box 176, SE-221 00, Lund, Sweden (N. Paracini).

## Abstract

Compositional asymmetry between lipid bilayer leaflets is a defining feature of biological membranes, yet its role in modulating protein binding remains largely unexplored. Here, we investigate how leaflet-specific composition affects the interaction between Myelin Basic Protein and biomimetic myelin membranes using neutron reflectometry. Asymmetric supported myelin bilayers containing deuterated cholesterol in the cytoplasmic leaflet enabled resolution of structural asymmetry. Neutron reflectometry measurements show that Myelin Basic Protein binds preferentially to asymmetric supported bilayers mimicking native myelin, whereas *Experimental Autoimmune Encephalomyelitis*-modified compositions exhibit weaker binding and more pronounced protein insertion. Disruption of asymmetry—either by thermal-induced lipid redistribution or by using symmetric cytoplasmic myelin—leads to a marked reduction in Myelin Basic Protein binding, despite increased membrane charge. Following vesicle adsorption and formation of a second bilayer, the protein layer narrows to near in vivo thickness in both systems, with the diseased condition exhibiting a wider inter-bilayer spacing. These results underscore the relevance of lipid asymmetry and composition in governing protein – membrane interactions, with implications for the molecular basis of myelin stability and demyelination.

## Introduction

Biological membranes play a central role in maintaining cellular integrity, enabling signal transduction, and regulating metabolic processes. A thorough understanding of the physicochemical properties of membrane constituents—and their interactions with both peripheral and integral membrane proteins—is essential to elucidate the molecular mechanisms underlying physiological function and disease. Among the various pathologies linked to altered membrane architecture, neurodegenerative disorders involving myelin sheath degradation stand out for their complexity and their significant impact on human health and disease.^1–3^

Myelin is a multilamellar lipid membrane that surrounds neuronal axons in the central and peripheral nervous systems, providing electrical insulation and allowing the rapid saltatory propagation of action potentials.^4–5^ It contains a diverse mixture of lipids, differing in headgroup chemistry, acyl chain length, and saturation. Cholesterol is also abundant, accounting for approximately 35 mol% of the membrane.^6^ The cytoplasmic and periplasmic leaflets (denoting the leaflets oriented toward the axoplasm and the extracellular apposition, respectively) of the myelin bilayer differ significantly in lipid composition, creating a pronounced asymmetry between both leaflets.^7^

The structure of myelin in vivo has long been the subject of extensive investigation, particularly through the use of X-ray techniques.^8–9^ These studies have demonstrated that, under proper cryoprotective conditions, the ultrastructure of myelin can be preserved with minimal alteration, allowing compacted and native-period membrane phases to coexist within the sheath. Purified myelin membranes from living organisms have also been widely studied using various model membrane approaches.^10–12^ For example, Langmuir monolayer experiments have revealed that under specific conditions, myelin lipids do not exist in a single lateral phase but rather segregate into at least two distinct phases: a more fluid and a more ordered one that predominantly retains cholesterol.^13–14^

In recent years, research has moved toward biomimetic systems that more accurately represent the lipid complexity and leaflet asymmetry of native myelin.^15–16^ These models have been employed to simulate both healthy and pathological states, for example by using lipid compositions derived from Experimental Autoimmune Encephalomyelitis (EAE), the standard animal model for Multiple Sclerosis (MS).

The myelin compaction and structural stability rely on the presence of Myelin Basic Protein (MBP), which anchors adjacent cytoplasmic leaflets of oligodendrocyte membranes, forming the dense membrane stacks characteristic of compact myelin.^17–18^ The interaction of MBP with model membranes has been the subject of numerous studies, particularly using liposomes and supported bilayers to mimic key aspects of myelin structure.^19–25^

Studies on MBP-membrane interactions have shown that the complex lipid composition critically affects membrane response. In particular, membranes with the diseased lipid profile exhibit instabilities upon MBP binding—such as loss of compact stacking—that resemble pathological features seen in demyelination.^26–28^ Our group has also previously demonstrated by small-angle x-ray and neutron scattering (SAXS and SANS) that while both native cytoplasmic and EAE-modified myelin vesicles tend to aggregate in the presence of MBP, only those with a native lipid composition progressively develop multilamellar stacks over time (aging).^29^ Moreover, MBP exhibited stronger interaction energy with cytoplasmic myelin membranes having the native composition as compared to the EAE-modified one, and native cytoplasmic myelin membranes had a higher bending stiffness than EAE-diseased membranes.^29–33^ Lee et al. showed by atomic force microscopy (AFM) that at low MBP concentrations, the protein preferentially accumulates in fluid membrane domains over more ordered ones.^16^ Taken together, these findings highlight the importance of lipid composition in modulating MBP binding and membrane mechanics.

The architecture of myelin represents an experimental challenge to decouple the roles of each leaflet and their interactions with MBP. In this context, the development of asymmetric supported myelin bilayers (SMB) provides a powerful platform for probing structure–function relationships at the molecular level. Although supported bilayers and vesicles with a symmetric profile have been widely studied in the past, only a limited number of studies have investigated asymmetric membranes^34–38^ or vesicles.^39^ However, membrane asymmetry is a hallmark of biological membranes, and its central importance is only slowly emerging.^40–42^ Knowledge on membrane asymmetry and its effects on membrane function or membrane-protein interactions is missing.

In the present work, a combination of Langmuir–Blodgett (LB) and Langmuir–Schaefer (LS) techniques were employed to fabricate planar asymmetric myelin membranes on silicon substrates. The incorporation of a deuterated lipid such as d₄₅-cholesterol in the cytoplasmic leaflet enabled contrast variation in neutron reflectometry (NR) experiments for resolving subtle differences in the bilayer composition. Asimilar approach was used in Clifton et al. ^35^ for studying the effect of asymmetry in a Gram-negative bacterial outer membrane biomimetic model.

NR was used to examine how the structure normal to the interface is modified when native and EAE-modified asymmetric myelin interact with MBP. The stability of the asymmetry was tested at different temperatures and compared with symmetric cytoplasmic myelin membranes in terms of the protein binding affinity. The out-of-plane structure was finally characterized when a second bilayer was deposited on the asymmetric SMB-MBP system.

These results provide new insights on how lipid complexity and membrane asymmetry modulate MBP-myelin interactions, offering a molecular perspective that is crucial to demyelination, human health and disease.

## Results and Discussion

To elucidate the interaction between MBP and SMB with high structural precision, we employed NR. This technique offers exceptional sensitivity to variations in the atomic composition normal to the membrane plane, enabling the reconstruction of depth-resolved SLD profiles with sub-nanometre resolution. Under specular reflection conditions—where the angle of incidence equals the angle of reflection—, the measured intensity provides direct insight into the stratification and composition of interfacial layers. This makes NR particularly well suited for probing the vertical architecture of model membrane systems, such as supported bilayers, and for assessing subtle structural rearrangements induced by protein binding.

### Validation of Membrane Asymmetry in SMB

To emulate CNS myelin, our model membranes comprise the major glycerophospholipids (phosphatidylcholine, phosphatidylethanolamine, phosphatidylserine), sphingolipids (sphingomyelin, cerebrosides, and sulfatides), and cholesterol, arranged asymmetrically between cytoplasmic and periplasmic leaflets as reported in Min et al. ^33^ (full molar profiles in Table S1, SI).

Asymmetric SMB with the native myelin lipid composition were initially assembled at 15 °C using a combination of the LB and LS techniques (see Experimental section at SI for details). The periplasmic leaflet was first deposited via the LB method to form the initial Langmuir monolayer, after which the bilayer was completed by transferring the cytoplasmic leaflet using the LS technique. In the adopted geometry, the periplasmic leaflet is oriented towards the silicon substrate, while the cytoplasmic leaflet interfaces with the bulk aqueous phase (see Fig. 1). NR measurements were conducted on two bilayer systems: one comprising a fully protiated cytoplasmic leaflet, and the other in which ovine cholesterol was selectively replaced with its deuterated analogue, d₄₅-cholesterol. The d₄₅-cholesterol was selected as the deuterated component because it was available with a well-defined deuteration level (∼79 ± 2%), allowing for accurate calculations of the SLD. Moreover, unlike phospholipids (used here as natural mixtures), cholesterol is a single, well-defined molecule, making selective deuteration straightforward. Asummary of the SLD values for all lipid and solvent components used in this study is provided in Table 1.

**Figure 1:**
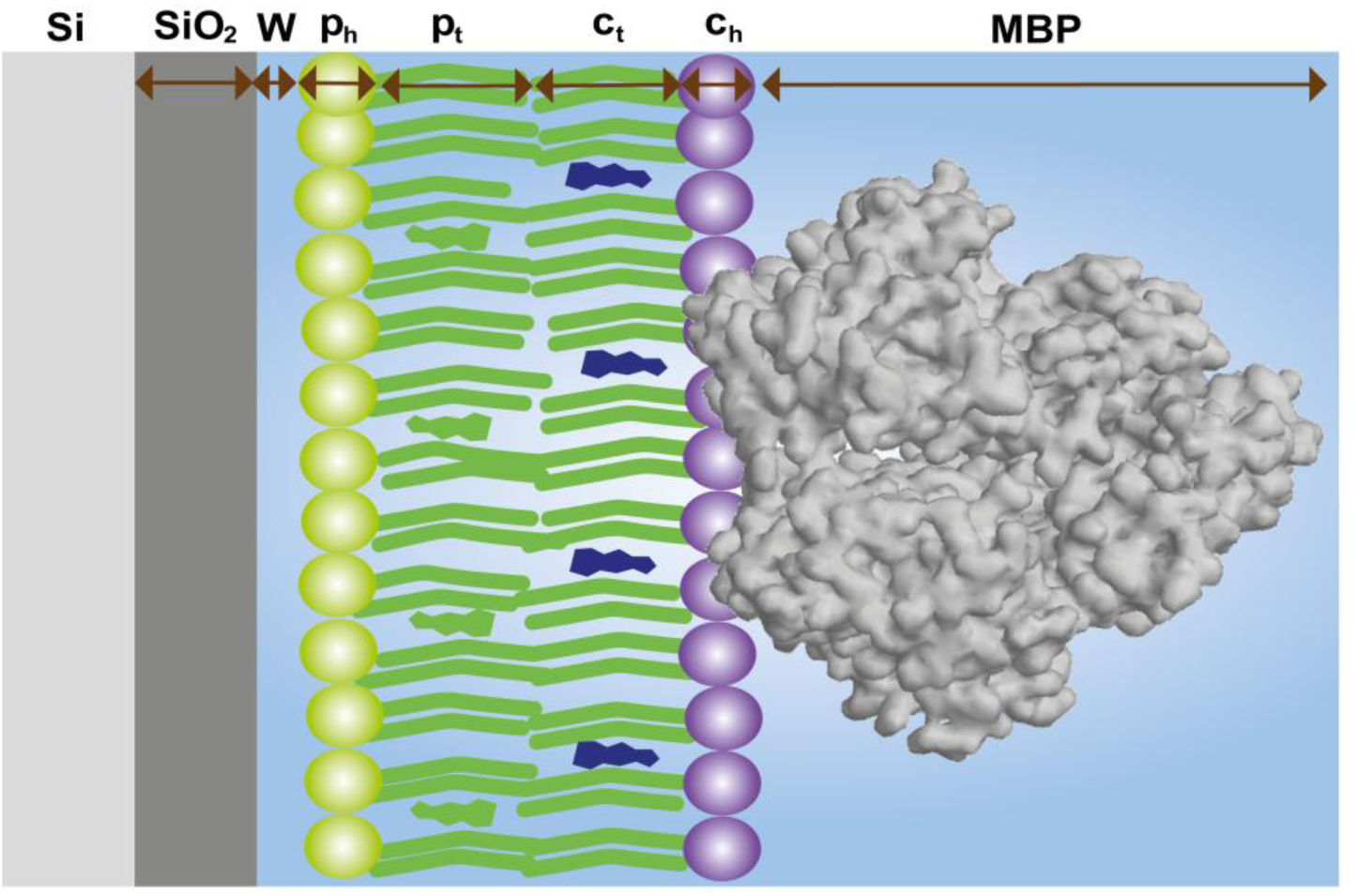
Schematic of the asymmetric SMB used in NR. From the substrate to the bulk: silicon (Si), silicon oxide (SiO₂), an interfacial water layer (W), inner periplasmic heads (*p_h_*) and tails (*p_t_*), and outer cytoplasmic tails (*c_t_*) and heads (*c_h_*). Where indicated in the manuscript, the cytoplasmic tail contains d₄₅-cholesterol (blue) to provide isotopic contrast. “Inner/outer” designate periplasmic/cytoplasmic sides relative to the substrate. Drawings are not to scale; colours denote model layers only.

**Table 1:** Theoretical neutron SLD (×10⁻⁶ Å⁻²) for buffers, substrate and membrane-associated layers used in the slab model at pH/pD 7.4. Values are fixed during fitting unless stated otherwise.

| Lipid/solvent | SLD ($\times 10^{-6} \text{ \AA}^{-2}$) |
| --- | --- |
| D<sub>2</sub>O MOPS buffer (pD 7.4) | 6.35 |
| 4MW MOPS buffer (pH 7.4) | 4.00 |
| SMW MOPS buffer (pH 7.4) | 2.07 |
| H<sub>2</sub>O MOPS buffer (pH 7.4) | -0.56 |
| Silicon | 2.07 |
| Silicon oxide | 3.41 |
| Native protiated $c_h$ | 1.79 |
| Native protiated $c_t$ | -0.09 |
| Native protiated $p_t$ | -0.21 |
| Native protiated $p_h$ | 1.92 |
| Native deuterated $c_t$ | 2.55 |
| EAE protiated $c_h$ | 1.76 |
| EAE protiated $c_t$ | -0.16 |
| EAE protiated $p_t$ | -0.04 |
| EAE protiated $p_h$ | 1.99 |
| EAE deuterated $c_t$ | 2.42 |

Each bilayer was characterized under four isotopic solvent contrasts —D₂O-buffer, four-matched water (4MW, 66% D₂O/34% H₂O), silicon-matched water (SMW, 38% D₂O/62% H₂O) and H₂O-buffer— yielding a total of eight distinct reflectivity profiles. Fig. 2 presents the resulting data, the corresponding model fits, and the derived SLD profiles for both the protiated and deuterated labelled bilayers.

**Figure 2:**
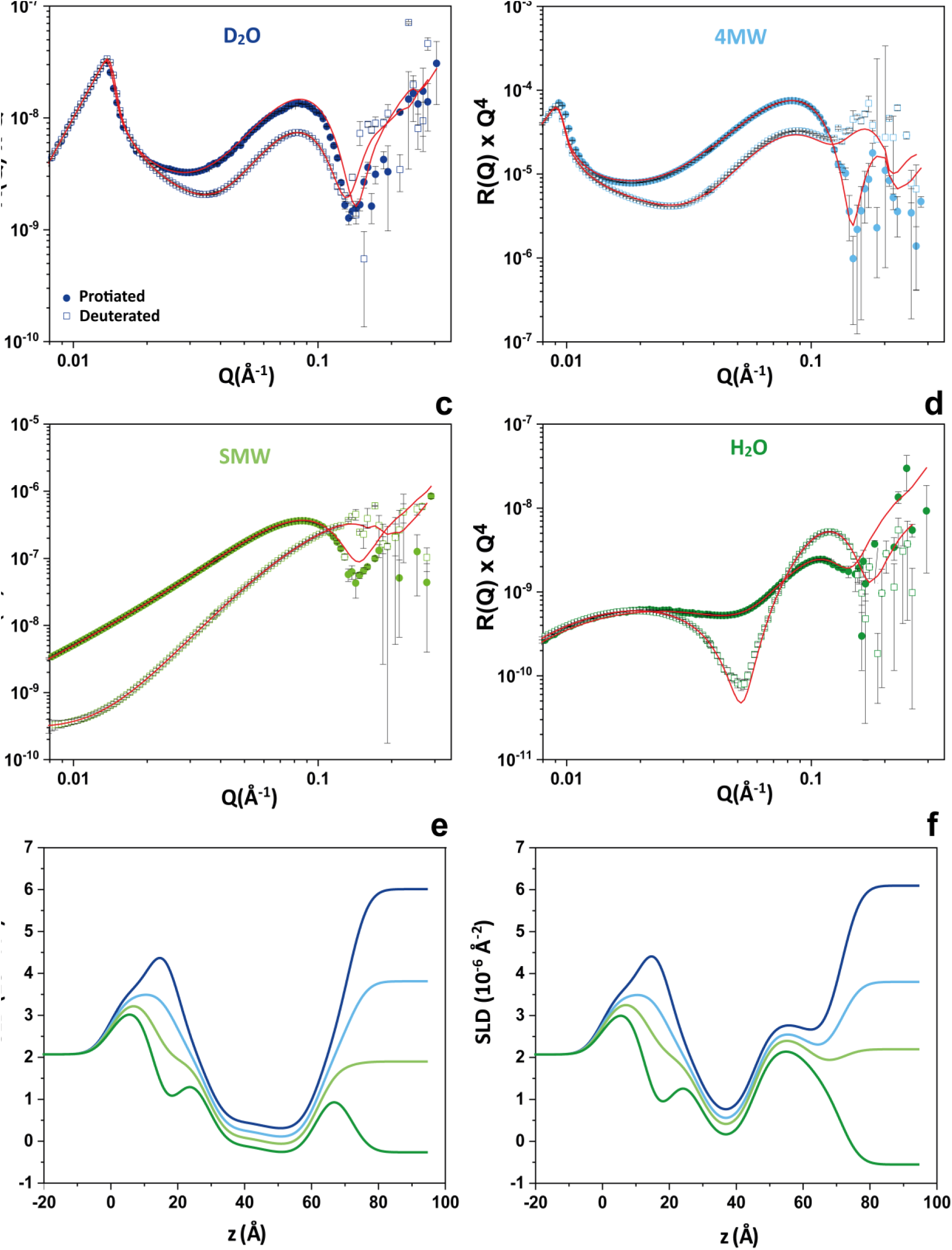
Neutron reflectivity (NR) and global co-refinement fits for a native asymmetric SMB with either a protiated or a deuterated cytoplasmic leaflet under four solvent contrasts: (a) D₂O, (b) 4MW, (c) SMW, and (d) H₂O. Symbols denote experimental data—filled circles: protiated, open squares: deuterated—and colours encode the solvent contrast (D₂O: dark blue; 4MW: light blue; SMW: olive; H₂O: green). Solid lines are the best co-refinement fits. Error bars represent 1σ. (e,f) Reconstructed SLD profiles (same colour coding).

The primary aim of these experiments was to assess whether membrane asymmetry is retained upon assembly at 15 °C. As summarised in Table 1, the SLD contrast between the acyl chain regions of the periplasmic and cytoplasmic (*c_t_* and *p_t_*) leaflets is negligible when both are protiated. In contrast, the introduction of deuterated cholesterol in the cytoplasmic leaflet produces a marked divergence in both the reflectivity curves and the derived SLD profiles as shown in Fig. 2, thereby providing evidence that asymmetry is maintained in native myelin membranes under these conditions. As expected from contrast variation, the non-deuterated bilayer (●) shows larger Kiessig fringes in D₂O and 4MW, whereas the deuterated one (□) dominates in H₂O, reflecting the reversal of ΔSLD with the solvent.

Using the six-layer model described in the Methods section, we fitted the eight contrasts per membrane. For the fitting procedure, all SLD values were held constant except for the SLD of the acyl chain regions, which was refined within a narrow range. From the hydration fraction *ϕ*_*w*_ we calculated a surface coverage fraction *η* defined as *η* = 1 − *ϕ*_*w*_. The adjusted structural parameters, including *η*, thickness (*t*) and SLD of the acyl chains, are summarised in Table 2.

**Table 2:**
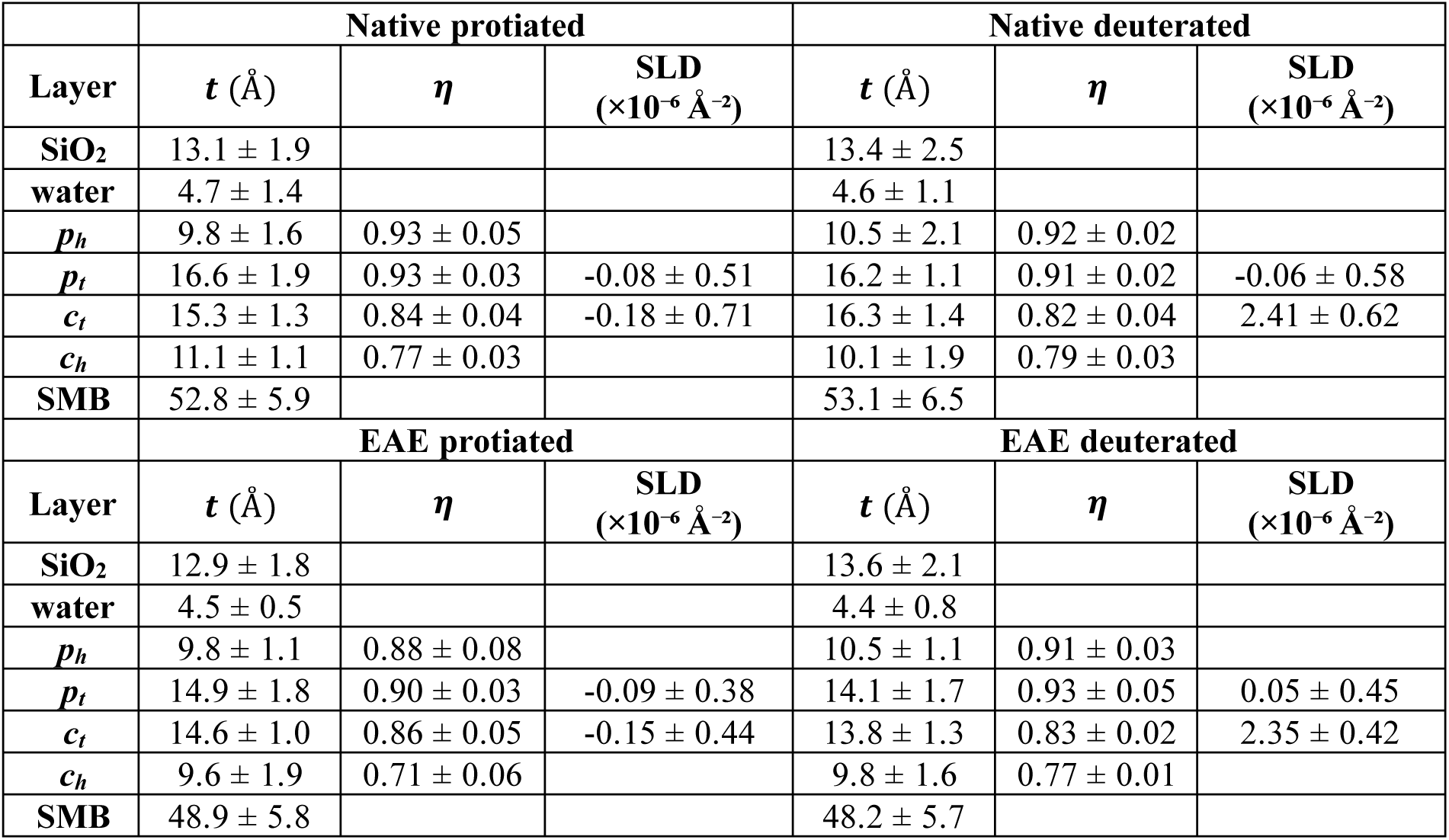
Fitted structural parameters for asymmetric SMB (native and EAE-modified) at 15 °C: layer thickness (*t*), surface coverage (*η* = 1 − *ϕ*_*w*_) and SLD (×10⁻⁶ Å⁻²). SMB denotes the total bilayer thickness (*p_h_* + *p_t_* + *c_t_* + *c_h_*). Uncertainties are 1σ from the fit covariance; for the totals, errors were obtained by covariance-aware propagation.

The resulting fits revealed high surface coverage for both bilayers. The periplasmic leaflet exhibited consistent coverage of 91–93%, regardless of cytoplasmic composition and the cytoplasmic leaflet showed slightly lower values, close to 80%. The *t* values obtained are in good agreement with expectations, yielding a total bilayer thickness around *t* = 53 Å, comparable to those reported for myelin lipids using SAXS and Brewster Angle Microscopy (BAM).^43^ Additionally, the thickness of the interfacial water layer between the silicon oxide and the bilayer ranged from *t* ∼ 4.5-5 Å.

To test whether the asymmetric assembly introduced any interleaflet exchange during LS deposition, we relaxed the usual constraint on the acyl chain SLD and fitted them with bounds around their theoretical values in the d-labelled bilayer. For the d-labelled bilayer the cytoplasmic-tail SLD was 2.41 ± 0.62 × 10⁻⁶ Å⁻² versus the theoretical 2.55 × 10⁻⁶ Å⁻² (≈ 5 % lower), while the periplasmic-tail SLD increased from −0.21 to −0.06 ± 0.58 × 10⁻⁶ Å⁻²; both shifts lie within uncertainty. Such minor offsets likely reflect natural variability in fatty-acyl composition and differences between assumed and effective molecular volumes, and/or slight mixing during LS transfer, rather than genuine interleaflet exchange at 15 °C. Importantly, no systematic drift in either leaflet’s SLD was detected over ∼9 h at 15 °C, arguing against cholesterol flip-flop under these conditions.

The same procedure was applied to the asymmetric EAE-modified myelin membrane. Reflectivity profiles were successfully collected for the four previously described contrasts (Fig. S1), yielding good fits across all eight conditions (protiated and deuterated cytoplasmic leaflets in each contrast). As with the native SMB, the structural values obtained were virtually identical between the deuterated and protiated versions, reinforcing the notion that d₄₅-cholesterol does not substantially alter the membrane’s structural properties.

Compared with native SMB (∼53 ± 6 Å), the EAE system was not significantly thinner (∼48 ± 6 Å) (Fig. 3; Table 2). To test for LS-induced interleaflet exchange, we slightly relaxed the usual acyl-chain SLD constraints; in the EAE d-labelled bilayer the cytoplasmic-tail SLD refined to 2.35 ± 0.42 × 10⁻⁶ Å⁻² versus the theoretical 2.42 × 10⁻⁶ Å⁻², i.e. within uncertainty. No systematic SLD drift was detected over ∼9 h at 15 °C.

**Figure 3:**
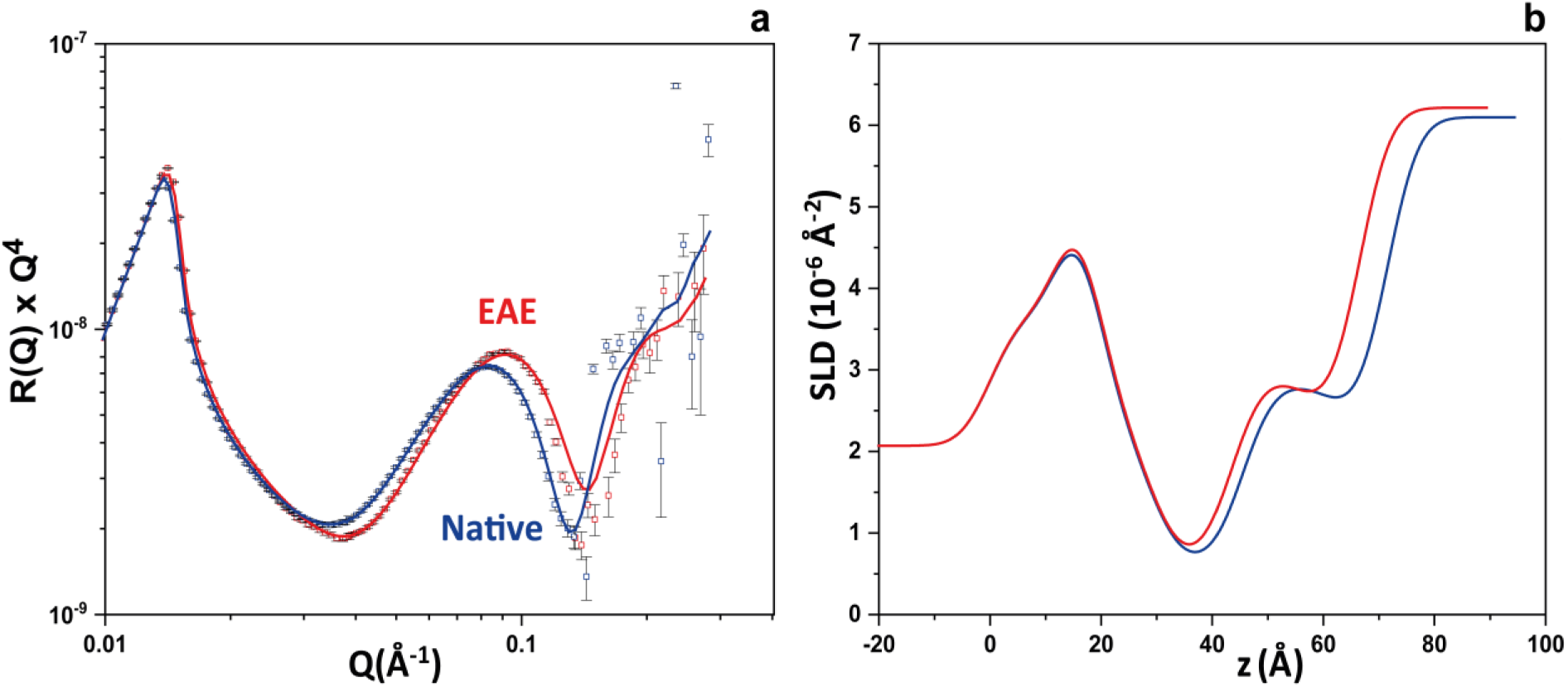
Comparison of NR (a) and corresponding SLD profiles (b) for native (blue) and EAE-modified (red) asymmetric SMB with a deuterated cytoplasmic leaflet in D₂O at 15 °C. Solid lines are co-refinement fits. Fitted parameters with 1σ uncertainties are reported in Table 2.

### Thermal Destabilisation of Membrane Asymmetry in SMB

Fig. 4a shows the reflectivity curves for a native asymmetric SMB featuring a deuterated outer leaflet, recorded at four temperatures between 15 and 37 °C, approaching physiological conditions. After each target temperature was reached, the sample was equilibrated for 15 minutes prior to measurement. Data were collected under two solvent contrasts (D₂O and H₂O-buffer), and comparative reflectivity profiles in D₂O are presented. Beyond a physicochemical benchmark, this series tests whether leaflet asymmetry can persist at 37 °C without active maintenance.

**Figure 4:**
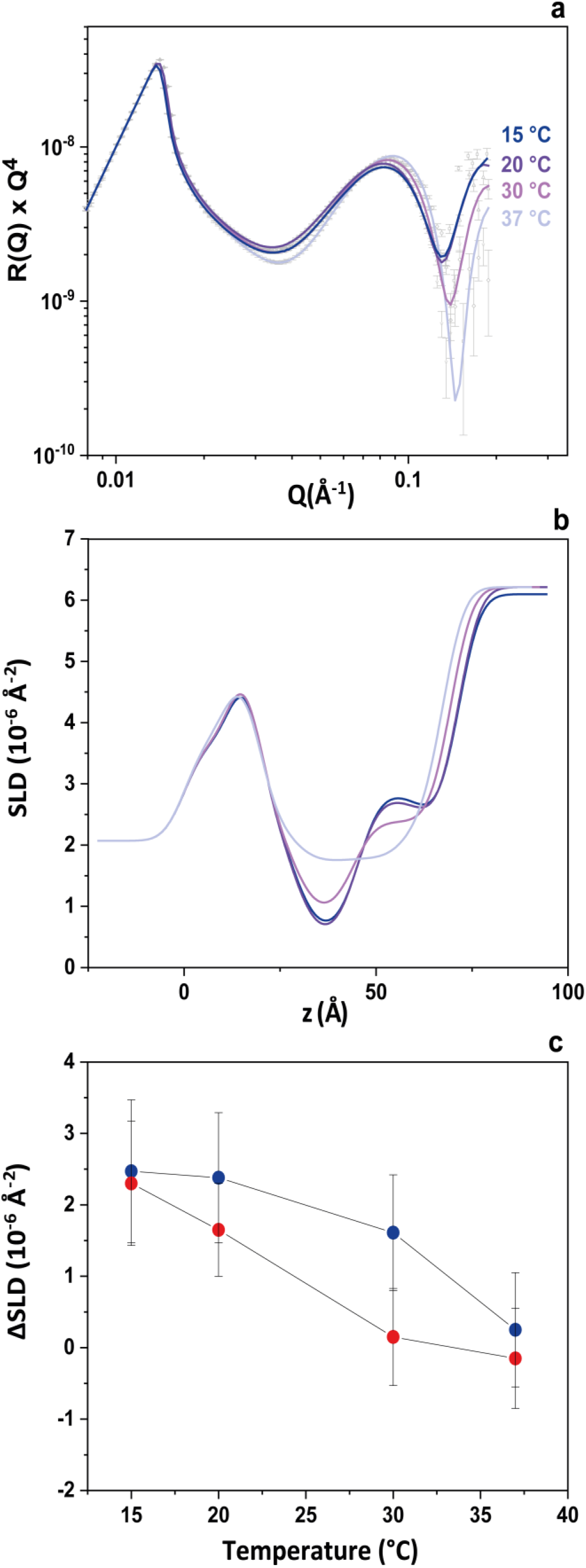
Temperature-dependent evolution of an asymmetric native SMB with a deuterated cytoplasmic leaflet: (a) reflectivity in D₂O at 15, 20, 30 and 37 °C; (b) SLD profiles; (c) acyl-chain SLD contrast between leaflets (ΔSLD = SLD*c_t_* − SLD*p_t_*) versus temperature for native (blue) and EAE-modified (red) SMB. Error bars in (c) are 1σ from the fit covariance with standard propagation.

Using the same six-layer interfacial model described earlier, the thickness and SLD of the acyl chain regions in both leaflets were refined to generate structural profiles of the bilayers at each temperature (Fig. 4b). To minimise the number of fitting parameters and based on the similarity in SLD values for the polar headgroup regions across the bilayer, both the SLD and *t* of these polar layers were fixed to the values reported in Table 1 and 2, respectively. Between 15 and 20 °C, structural changes are within error. With increasing temperature, the acyl chain region thins modestly (about 4 Å from 15 to 37 °C; Table S2), while the leaflet SLD contrast (ΔSLD = SLD*c_t_* − SLD*p_t_*) decreases significantly from 15 to 37 °C for both compositions, indicating cholesterol redistribution (Fig. 4c; Table 3). The EAE-modified SMB shows an earlier loss of contrast, with ΔSLD already near zero at 30 °C, whereas the native system decays more gradually (Fig. 4c; Table 3). Changes at intermediate temperatures are small and, within the propagated uncertainties, do not resolve composition-specific differences; we therefore focus on the robust 15 → 37 °C decrease. EAE-modified counterparts of panels (a) and (b) are provided in the Supporting Information (Fig. S2).

**Table 3:** Temperature series of leaflet SLD and contrast (ΔSLD = SLD*c_t_* − SLD*p_t_*). Leaflet acyl-chain SLD for the periplasmic (*p_t_*) and cytoplasmic (*c_t_*) leaflets and their contrast ΔSLD at 15, 20, 30 and 37 °C for native and EAE-modified asymmetric SMB. Uncertainties are 1σ from the fit covariance; Δ-type quantities use full covariance propagation.

| $T$ (°C) | SLD ($\times 10^{-6} \text{ \AA}^{-2}$) | | | | | |
| --- | --- | --- | --- | --- | --- | --- |
|  | Native |  |  | EAE-modified |  |  |
| | $p_t$ | $c_t$ | $c_t - p_t$ | $p_t$ | $c_t$ | $c_t - p_t$ |
| 15 °C | -0.06±0.38 | -2.41±0.62 | 2.47±1.00 | 0.05±0.45 | 2.35±0.42 | 2.30±0.87 |
| 20 °C | 0.01±0.47 | 2.39±0.44 | 2.38±0.91 | 0.40±0.36 | 2.05±0.29 | 1.65±0.65 |
| 30 °C | 0.41±0.39 | 2.02±0.42 | 1.61±0.81 | 0.90±0.26 | 1.05±0.42 | 0.15±0.68 |
| 37 °C | 1.11±0.43 | 1.36±0.37 | 0.25±0.80 | 1.10±0.39 | 0.95±0.31 | -0.15±0.70 |

This interpretation is in line with previous work by Gerelli et al.^38^, who reported that appreciable interleaflet mixing in asymmetrically deposited phospholipid bilayers occurs predominantly when the lipids are in the fluid phase. A similar temperature dependence has been observed for cholesterol flip-flop, which accelerates markedly above 17 °C and occurs on a sub-millisecond timescale near physiological temperatures (∼37 °C), as demonstrated by Gu et al. ^44^ using atomistic molecular dynamics simulations.

This experiment also demonstrates that in the absence of protein components, the lipid asymmetry preserved in myelin *in vivo* at physiological temperature is not maintained in the SMB. This highlights the significant contribution of additional biological factors such as lipid transport proteins, cytoskeletal anchoring, and interactions with the extracellular matrix in maintaining membrane asymmetry *in vivo.*^45–46^

### Differential MBP Binding to Native and EAE-modified Asymmetric SMB

As a next step, MBP was introduced into the liquid cell at a concentration of *c*_*MBP*_ = 0.1 mg/mL in the buffer solution for both the native and EAE-modified asymmetric SMB with the d-cytoplasmic outer leaflet. The experiments were conducted at 15 °C. Owing to electrostatic attraction, MBP interacts with the negatively charged membrane surface, forming a protein-rich layer atop the bilayer. Reflectivity data in D₂O for both systems in the presence of MBP, along with their corresponding SLD profiles and schematic illustrations of the membrane architecture, are shown in Fig. 5.

**Figure 5:**
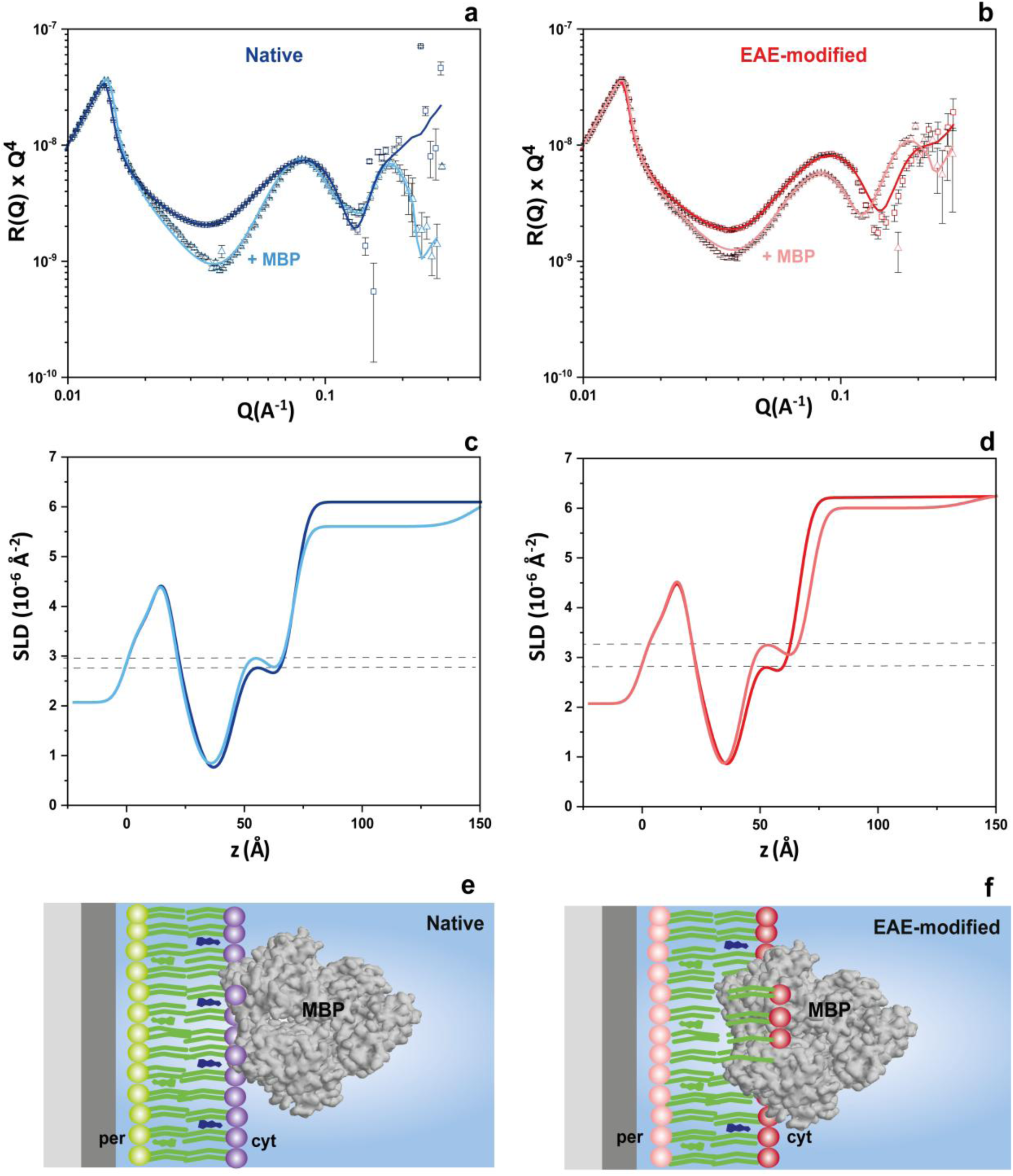
NR data (a,b), SLD profiles (c,d), and schematics (e,f) for native (blue) and EAE-modified (red) asymmetric SMB before and after MBP addition at 15 °C (D₂O). MBP thickness *t*_*MBP*_ and coverage *η*_*MBP*_ are summarised in Table 4.

**Table 4:** Fitted parameters for native and EAE-modified asymmetric SMB in the presence of MBP at 15 °C: layer thicknesses (*t*), surface coverage (*η*), and SLD (×10⁻⁶ Å⁻²) for the SMB, and *t*_*MBP*_ with *η*_*MBP*_ for the protein. Values are best-fit ±1σ.

|  | Native |  |  | EAE-modified |  |  |
| --- | --- | --- | --- | --- | --- | --- |
| Layer | $t \text{ (\AA)}$ | $\eta$ | SLD<br>($10^{-6} \text{ \AA}^{-2}$) | $t \text{ (\AA)}$ | $\eta$ | SLD<br>($10^{-6} \text{ \AA}^{-2}$) |
| $p_h$ | $10.1 \pm 1.5$ | $0.92 \pm 0.02$ | $1.96 \pm 0.61$ | $10.0 \pm 1.1$ | $0.89 \pm 0.03$ | $1.95 \pm 0.48$ |
| $p_t$ | $16.2 \pm 1.3$ | $0.91 \pm 0.02$ | $-0.08 \pm 0.51$ | $14.5 \pm 2.1$ | $0.90 \pm 0.06$ | $0.19 \pm 0.71$ |
| $c_t$ | $17.3 \pm 1.4$ | $0.79 \pm 0.04$ | $2.47 \pm 0.44$ | $17.5 \pm 1.6$ | $0.77 \pm 0.04$ | $3.01 \pm 0.88$ |
| $c_h$ | $10.3 \pm 1.5$ | $0.75 \pm 0.03$ | $1.71 \pm 0.52$ | $11.5 \pm 1.2$ | $0.75 \pm 0.05$ | $2.01 \pm 0.73$ |
| **SMB** | $53.9 \pm 5.7$ | | | $53.5 \pm 6.0$ | | |
| **MBP** | $78.9 \pm 8.9$ | $0.25 \pm 0.04$ | | $68.6 \pm 9.1$ | $0.10 \pm 0.04$ | |

These datasets were modelled using a sequential slab structure comprising water -*p_h_* -*p_t_*-*c_t_ -c_h_*, with an additional MBP layer positioned on the solvent-facing side. The MBP layer was fitted with two free parameters: its thickness *t*_*MBP*_ and its coverage *η*_*MBP*_. The underlying SMB parameters were initialised from the protein-free fits (Table 2) and relaxed according to the constraints defined in NR data analysis (SI); results are summarised in Table 4.

The *R*(*Q*) × *Q*⁴ plots (Fig. 5a,b) show composition-dependent responses to MBP: in native SMB the minima remain essentially fixed in *Q*, whereas in EAE they shift to lower *Q* and SLD fits indicate bilayer thickening; accordingly, we analyze the two datasets separately. For native myelin, the model yielded a well-defined MBP layer with *t*_*MBP*_ ≈ 80 Å and a surface concentration (defined as *c*_*MBP*/*SMB*_ (%) = *η*_*MBP*_ ∗ 100) of *c*_*MBP*/*SMB*_ = 25 ± 4 % vol. The structural parameters of the underlying SMB remained unchanged compared to the protein-free system (*t* = ∼53.1 ± 6.5 Å → 53.9 ± 5.7 Å and SLD*c_t_* = 2.41 ± 0.62 → 2.47 ± 0.44 × 10⁻⁶ Å⁻², respectively), suggesting minimal insertion of MBP into the bilayer (Table 4 and Fig. 5d). This result aligns with previous NR data on recombinant MBP, which showed a similarly thick protein layer with ∼25% volume occupancy atop model bilayers, supporting a conserved surface binding mode under native-like conditions.^31^

In contrast, some differences were observed for the EAE-modified asymmetric SMB. The protein layer was thinner by approximately 10 Å, and the surface enrichment was lower, with *c*_*MBP*/*SMB*_ = 10 ± 4 % vol (Fig. 6a). Additionally, the total thickness and SLD of the outer leaflet with MBP increased relative to the bare SMB (*t* = ∼48.2 ± 5.7 Å → 53.5 ± 6.0 Å and SLD*c_t_*= 2.35 ± 0.42 → 3.01 ± 0.88 × 10⁻⁶ Å⁻², respectively), indicating a more pronounced protein penetration into this leaflet in the EAE-modified system (Fig. 5d). This differential adsorption is also in line with previous observations showing that MBP binding is strongly modulated by membrane composition and domain organization, with reduced affinity on EAE-like lipid environments.^16^

**Figure 6:**
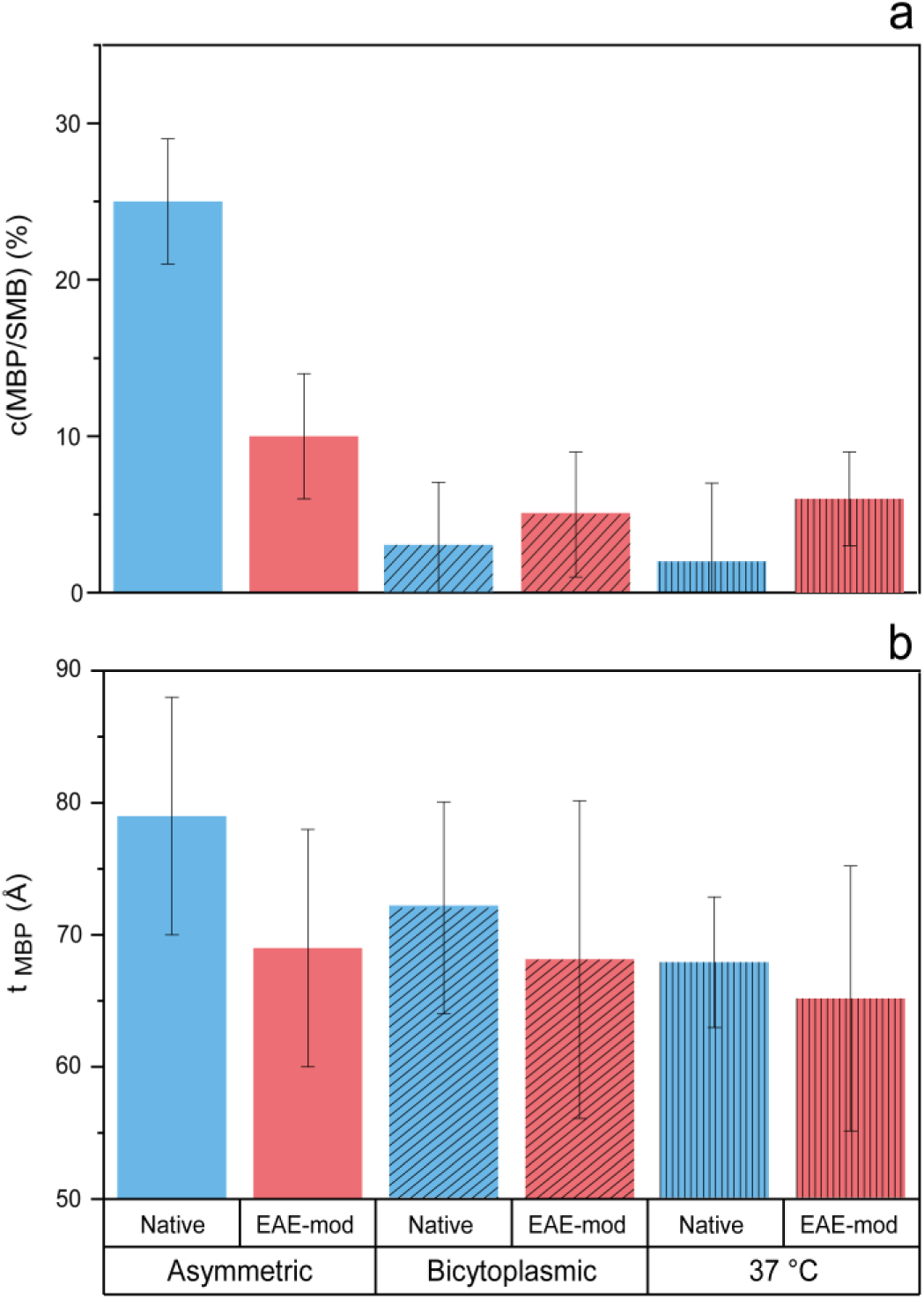
MBP binding across membrane states—asymmetric (15 °C), symmetric bicytoplasmic (15 °C), and asymmetric heated (37 °C). (a) Surface concentration *c*_*MBP*/*SMB*_ (%) (b) Protein-layer thickness *t*_*MBP*_. Blue bars: native; red bars: EAE-modified. Bars show best-fit values; error bars are 1σ. Exact values in Tables 4 and S3.

### Reduced MBP Binding Upon Loss of Membrane Asymmetry

The interaction between MBP and SMB was further investigated under two additional conditions. First, a native symmetric SMB was assembled by depositing two deuterated cytoplasmic leaflets at 15 °C. In the absence of MBP, this symmetric SMB matched the asymmetric SMB at the same temperature in overall thickness (Table S3) and outer-leaflet composition; nevertheless, *c*_*MBP*/*SMB*_ (%) changed substantially. Although the bicytoplasmic bilayer contains a higher overall PS fraction, the MBP-facing leaflet has the same PS content as in the asymmetric SMB. The markedly lower *c*_*MBP*/*SMB*_ (∼3 vol %; Fig. 6) therefore cannot be attributed to a reduced interfacial charge but rather to the loss of leaflet asymmetry. This value is consistent with the one reported by Krugmann et al.^30^ for cytoplasmic myelin bilayers formed via vesicle fusion and is clearly lower than the 25 % vol for the asymmetric native SMB (vide supra). A similar trend with less pronounced reduction was observed for the EAE-modified composition, where *c*_*MBP*/*SMB*_ decreased from 10% to about 5% vol between the asymmetric and the bicytoplasmic SMB. Reflectivity and SLD profiles remained mostly unchanged upon MBP addition (Fig. S3 and Table S3), consistent with negligible bilayer perturbation.

The asymmetric SMB was also heated to 37 °C, a temperature at which cholesterol redistribution across the bilayer was shown to occur, leading to a symmetric SMB. Under this condition, MBP binding was further reduced in both the native and EAE-modified compositions. Specifically, *c*_*MBP*/*SMB*_ decreased drastically from 25% vol to ∼2% vol in the native myelin and from 10% vol to ∼6% vol in the EAE-modified case (Fig. 6a), reaching values comparable to those of the symmetric bicytoplasmic membranes. Addition of MBP caused only minor variations in the reflectivity and SLD profiles at 37 °C (Fig. S3 and Table S3). Together, these results demonstrate that the affinity of MBP for myelin is markedly reduced when the membrane loses its leaflet compositional asymmetry irrespective of the mechanism leading to a gain of membrane symmetry.

A clear thinning of the MBP layer (Fig. 6b) in the EAE-modified membrane compared to the native one is only apparent for the asymmetric SMB—the condition also showing the strongest MBP binding. For both the symmetric bicytoplasmic bilayer and the asymmetric SMB rendered symmetric upon heating to 37 °C, the *t*_*MBP*_ values for native and EAE-modified compositions remain essentially unchanged within the propagated uncertainties.

### Vesicle Adsorption Forming a Double Bilayer (DBL)

To probe whether MBP-mediated adhesion persists upon bringing a second membrane into contact, we deposited cytoplasmic myelin vesicles onto pre-formed MBP/SMB (native or EAE) and promoted rupture/coverage by an osmotic D₂O shock. This system can be considered as a simple biomimetic model of an oligodendrocyte wrapping around an axon, with the cytoplasmic membrane leaflets held together by MBP ^31^.

After ∼1 h, the reflectivity stabilised (Fig. 7a). In the raw *R*(*Q*) curves, vesicle deposition shifts the Kiessig minima to lower *Q* and reduces their spacing, consistent with the addition of a ∼50 Å slab and an increase in interfacial roughness; the effect is captured by the reconstructed SLD profiles (Fig. 7b).

**Figure 7:**
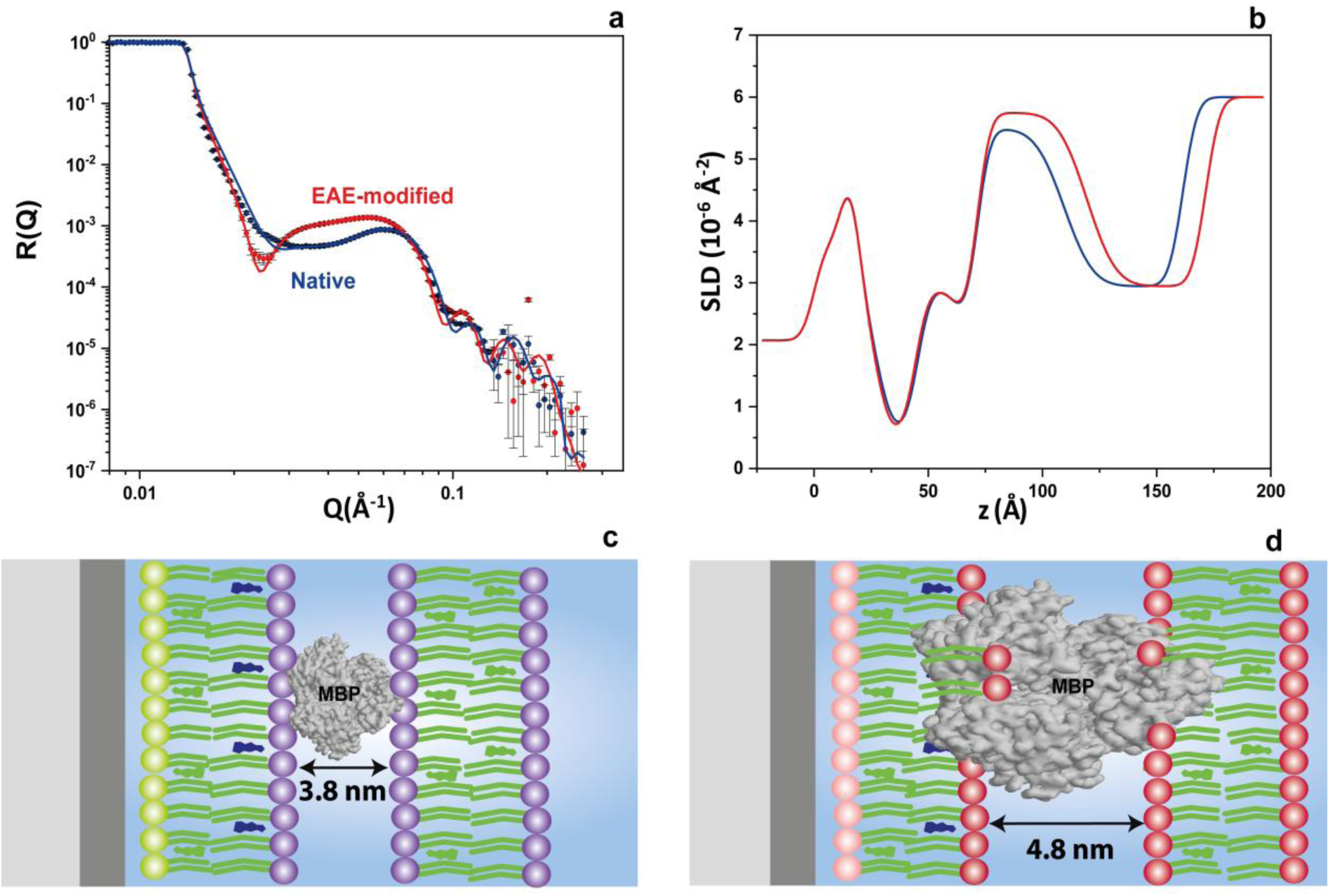
Double-bilayer (DBL) assembly on pre-adsorbed MBP/SMB: (a) NR data and fits; (b) SLD profiles; (c,d) schematics for native (blue) and EAE-modified (red) compositions. The DBL is modelled as a single, rough slab; MBP compacts upon bridging. Fitted MBP and DBL parameters are listed in Table 5.

**Table 5:**
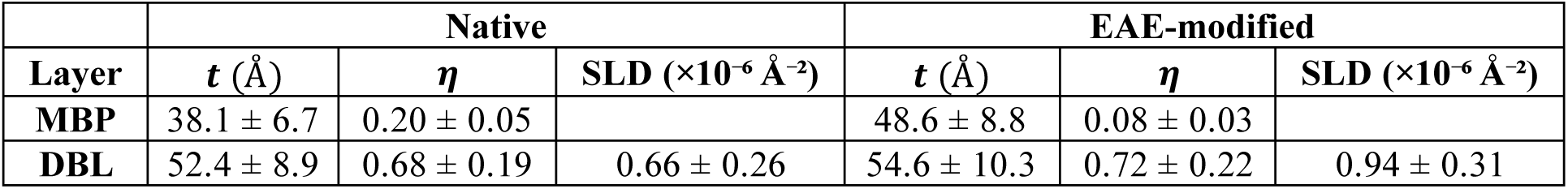
Fitted parameters for the MBP layer and the upper bilayer in the DBL configuration: MBP thickness (*t*_*MBP*/*DBL*_) and coverage (*η*_*MBP*/*DBL*_), and DBL apparent thickness (*t*_*DBL*_), coverage (*η*_*DBL*_) and SLD (×10⁻⁶ Å⁻²). Values are best-fit ±1σ.

Datasets were analysed with the DBL model defined in the Methods section: the underlying SMB (from the MBP/SMB stage) was kept fixed, the MBP volume fraction was allowed to vary within ±20 % around its pre-adsorbed value, and the upper bilayer was represented by a single, highly rough slab with adjustable SLD and thickness. Under these constraints, the MBP layer compacted to *t*_*MBP*/*DBL*_ = 38.1 ± 6.7 Å (native) and 48.6 ± 8.8 Å (EAE), while its coverage decreased modestly relative to the MBP/SMB state from *η* = 0.25 ± 0.04 to 0.20 ± 0.05 in native, and from *η* = 0.10 ± 0.04 to 0.08 ± 0.03 in EAE-modified myelin (Tables 4–5). This behaviour indicates a re-arrangement into a denser adhesive layer as MBP bridges the two cytoplasmic leaflets.

The upper bilayer is incomplete and laterally disordered: the single-slab returns an apparent coverage *η*_*DBL*_ of 0.68 ± 0.19 (native) and 0.72 ± 0.22 (EAE), and SLD of 0.66 ± 0.26 and 0.94 ± 0.31 × 10⁻⁶ Å⁻², respectively (Table 5). The higher SLD in the EAE case is compatible with deeper MBP penetration and/or enhanced D₂O uptake at the protein–membrane interface; given the single-slab representation, these effects cannot be disentangled unambiguously. Finally, the inter-bilayer spacing, proxied by *t*_*MBP*/*DBL*_, is on average ∼10 Å larger in EAE than in native (*Δt* = 10.5 ± 11.1 Å), whereas the native membranes consistently retain a higher MBP volume fraction. Together, the DBL experiment shows that MBP bridging compacts upon membrane–membrane contact, yet lipid composition chiefly tunes protein recruitment, even in the presence of an upper bilayer (Fig. 7c,d; Table 5).

## Conclusions

To our knowledge, this is the first study focused on the design of asymmetric membranes with realistic myelin-like lipid complexity, providing a new platform for dissecting leaflet-specific contributions to membrane–protein interactions. Our experiments demonstrate that membrane asymmetry is not a neutral feature in the organization of myelin bilayers, but rather a determining factor that modulates the MBP-membrane interactions.

Using NR, we showed that MBP binds preferentially to asymmetric SMB that mimic the native composition of myelin, whereas symmetric bicytoplasmic bilayers, despite bearing a higher net negative charge, exhibit significantly reduced protein binding. The reduced MBP recruitment to a bicytoplasmic membrane—despite comparable surface charge at the contacting leaflet—supports a mechanism in which leaflet asymmetry and interleaflet coupling tune the local packing and hydration landscape sensed by MBP; we therefore hypothesize that asymmetry pre-organizes an anionic, weakly hydrated cytoplasmic environment that enhances MBP binding. Previous studies using both simple and more complex membrane systems had already shown that changes in compositional asymmetry can influence the mechanical properties of membranes ^47–48^ and their interactions with proteins.^40, 49^

As the system reaches physiological temperature, the distribution of cholesterol across the leaflets notably shifts. Both spontaneous lipid redistribution at 37 °C and symmetric bicytoplasmic bilayers led to a marked decrease in MBP surface coverage, revealing the critical role of dynamic interleaflet coupling in maintaining membrane functionality. Upon heating pre-adsorbed samples to 37 °C, the MBP surface concentration decreased (native: ∼25 → ∼2 vol %; EAE: ∼10 → ∼6 vol %; Fig. 6), approaching the low values observed for bicytoplasmic bilayers. Asymmetric lipid distribution is known to be actively sustained in vivo by the coordinated action of flippases, floppases, and cytoskeletal scaffolds, often at considerable energetic cost.^45–46^ Our results suggest that disruption of this tuned equilibrium potentially triggered by oxidative stress or inflammatory cues *in vivo* may lead to scrambling, the non-selective, bidirectional redistribution of lipids across the bilayer, and ultimately alter the membrane architecture and signalling dynamics.^50^ In this context, the observed reduction in MBP binding affinity upon asymmetric perturbations may represent an early consequence of lipid scrambling, contributing later to membrane destabilization processes relevant under neuroinflammatory conditions. Given MBP’s capacity to bridge cytoplasmic leaflets and condense anionic lipids, it is possible that protein binding itself help stabilize leaflet asymmetry in vivo by hindering interleaflet lipid exchange.

In line with the results of Raasakka et al.^31^ and Krugmann et al.^29–30^, we found that the initial MBP layer thickness reaches ∼80 Å, more than twice the typical spacing of the intraperiod dense line reported for native myelin. This could reflect the extension of a single disordered MBP molecule or the formation of an amorphous protein meshwork with lateral self-association,^31, 51–52^ possibly involving dimers. It was also observed that, while MBP inserts more deeply into the modified membranes, it does not promote the same level of protein enrichment as in the native system. This points to an uncoupling between membrane insertion and adhesive function. Complementary DBL experiments provided a mechanical and biological perspective: after deposition of the second bilayer, the pre-adsorbed MBP layer became reproducibly thinner, indicating a rearrangement into a more compact and adhesive conformation as the protein bridges both leaflets. A key finding is that this transition appears less effective in the EAE model, where the intermembrane spacing remained consistently about 1 nm larger than in the native case. Altogether, these results support the idea that both MBP recruitment and structural adaptation are strongly influenced by the lipid environment, as previously proposed using similar biomimetic systems.^28–29^

In summary, these findings support the view that myelin is not a passive lipid sheath, but a dynamic structure whose stability relies on a tightly regulated interplay between protein conformation and lipid organization.^17, 53^ By providing nanoscale insight into MBP–membrane interactions under asymmetric and pathological conditions, our study offers mechanistic understanding of how membrane architecture governs protein function and, ultimately, myelin stability.

## Experimental Section

A complete description of experimental procedures is provided in the Supporting Information. Below we summarize the NR data analysis used to model stratified layers and co-refine multiple isotopic contrasts.

### NR Data Analysis

Reflectivity profiles were collected from a series of chemically equivalent myelin bilayer samples differing only in the isotopic (deuterium) labelling of the cytoplasmic leaflet. Specifically, the non-labelled cytoplasmic leaflet contained protiated cholesterol and the other labelled cytoplasmic leaflet its deuterated analogue d₄₅-cholesterol, while the periplasmic leaflet composition remained unchanged in both cases.

The base SMB was modelled with six layers (Fig. 1): native silicon oxide, interfacial water, periplasmic heads and tails (*p_h_*and *p_t_*), cytoplasmic tails and heads (*c_t_* and *c_h_*). Upon MBP addition, a seventh layer was introduced to represent the adsorbed protein, and in the final configuration with vesicle adsorption, an eighth layer accounting for the second bilayer was incorporated.

Data from FIGARO were processed using COSMOS^54^ within the LAMP suite (ILL), while data from INTER were reduced using the Mantid framework (ISIS). Data fitting was performed using the Anaklasis package ^55^, which allows co-refinement of multiple contrasts using a stratified layer model. A global scaling factor was introduced to account for minor attenuation effects and potential diffuse scattering, particularly in late-stage samples containing vesicles. For each dataset, layer thicknesses (*t*) and hydration fractions (defined as solvent volume fraction, *ϕ*_*w*_) were fitted within physically reasonable ranges. To resolve leaflet asymmetry in selectively deuterated samples, the acyl chain SLD were fitted independently for each leaflet. Headgroup SLD were kept at their theoretical values, while head and tail thicknesses were allowed to vary independently for inner and outer leaflets (heads: 7–12 Å; tails: 12–17 Å). Hydration was leaflet-specific (heads: 0–0.7; tails: 0–0.5). A single lipid–lipid roughness of *σ* = 3.0 ± 0.5 Å was applied; Si/SiO₂ and MBP roughnesses were fitted separately and kept within *σ* = 0–4 Å and 0–15 Å, respectively. The SiO₂ thickness and the interfacial-water gap were refined per dataset with bounds *t* = 9-15 and 2–10 Å, respectively. For d-cytoplasmic samples, the outer-tail SLD was allowed between 1.6 and 3.0 × 10⁻⁶ Å⁻², and the inner-tail SLD between −0.5 and 0.5 × 10⁻⁶ Å⁻² (broadened to −0.5–1.8 × 10⁻⁶ Å⁻² for the 37 °C series).

For datasets recorded after MBP addition, the structural parameters of the underlying SMB were initialised from the protein-free best fits (Table 2) and then allowed to relax to accommodate potential protein insertion. Unless noted otherwise, thickness, hydration and SLD for each leaflet were permitted to vary by up to ±20 % relative to their protein-free values. MBP SLD was computed from sequence using Biomolecular SLD calculator ^56^ assuming 90 % exchange of labile hydrogens, and MBP thickness was fitted within *t* = 60–100 Å.

Finally, liposomes were injected to the SMB/MBP coated wafers to form double bilayers (DBL) which were also measured in D_2_O. For DBL fits, the SMB obtained at the MBP/SMB stage was kept fixed, and the MBP volume fraction was allowed to vary by ±20 % around the pre-adsorbed value. The second bilayer was modelled as a single homogeneous slab with adjustable SLD between 0.3 and 0.9 × 10⁻⁶ Å⁻², a thickness *t* between 45 and 55 Å, and a high roughness (*σ* = 0-7 Å), in order to minimize the number of free parameters.

Unless noted otherwise, reported uncertainties are 1σ (68%) from the covariance matrix of the Levenberg–Marquardt minimizer in Anaklasis, with fits weighted by pointwise experimental errors and after convolution with the instrumental resolution appropriate to each dataset. For derived quantities (e.g., total bilayer and acyl region thicknesses and their temperature differences), we propagated the full joint covariance (i.e., including parameter correlations).

## Associated Content

### Supporting Information

It includes an experimental section (PDF), additional NR datasets and global co-refinement fits for EAE-modified asymmetric SMB (PDF), temperature-series NR and SLD profiles for native and EAE SMB; full thickness tables (PDF) and NR data and co-refinement fits for bicytoplasmic and cholesterol-redistributed SMB at 15/37 °C before/after MBP adsorption (PDF).

### Data Availability

The raw neutron reflectometry datasets supporting this study are openly available at the ISIS Neutron and Muon Source and the Institut Laue-Langevin data portals under the following DOIs: *10.5286/ISIS.E.RB2220200-1, 10.5286/ISIS.E.RB2410499-1*, and *10.5291/ILL-DATA.8-02-993*. Processed data and model-fit outputs are provided in the Supporting Information (PDF). Additional files (reduced reflectivity profiles and co-refinement input files) are available from the corresponding authors upon request.

## Author Information

### Author Contributions

J.M.P. and A.M.S. contributed equally to this work.

### Notes

The authors declare no competing financial interest.

## Acknowledgements

We gratefully acknowledge beam-time allocation for the neutron scattering experiments on the neutron reflectometers INTER at the ISIS Neutron and Muon Source (Didcot, UK) and FIGARO at the Institut Laue-Langevin (Grenoble, France). We also acknowledge the support of the Australian Government in provision of access to ANSTO’s National Deuteration Facility which is partly funded through the National Collaborative Research Infrastructure Strategy (NCRIS). We further acknowledge support from the Tasso Springer Fellowship Program.

