## Supplementary Information for "Myelin Basic Protein Binding Is Modulated by Leaflet Asymmetry and Lipid Composition"

##### The file includes:

- Experimental section
- Figures S1 to S3 and tables S1 to S3
- References

### Experimental Section

#### Materials

The biomimetic myelin membranes employed in this study were composed of porcine brain-derived lipids—namely phosphatidylcholine (PC), phosphatidylethanolamine (PE), phosphatidylserine (PS), sphingomyelin (SM), cerebrosides, and sulfatides—together with ovine cholesterol. All lipid species were sourced from Avanti Polar Lipids (Alabaster, AL, USA). As these lipids are extracted from natural tissues, they inherently possess a heterogeneous distribution of fatty acyl chains; the detailed fatty acid profiles are provided by the manufacturer on their website. In bilayers with a deuterated cytoplasmic leaflet, ovine cholesterol was replaced by its perdeuterated analogue, $d_{45}$-cholesterol (79 ± 2% of D enrichment), sourced from the National Deuteration Facility (NDF, ANSTO, Lucas Heights, NSW, Australia).

3-(N-morpholino)propanesulfonic acid (MOPS; 99%, Alfa Aesar, CAS 1132-61-2; cat. no. A12914) was used to prepare 10 mM $D_2O$ or $H_2O$ – buffer (pD /pH = 7.4). Myelin basic protein (MBP; bovine, purified; Sigma-Aldrich, cat. 13-104) was used as supplied, stored at −20 °C, and diluted in $D_2O$ or $H_2O$ – MOPS buffer (pD /pH = 7.4) immediately before experiments to a final concentration of $c_{MBP}$ = 0.10 mg $mL^{-1}$.

#### Lipid Mixture Preparation

Individual lipids were pre-dissolved in high-purity solvents (chloroform or chloroform:methanol 2:1 v/v) purchased from Sigma-Aldrich (≥99.5%, Sigma-Aldrich, St. Louis, MO, USA) and mixed at appropriate molar ratios to yield final concentrations of 1 mg/mL for Langmuir monolayer deposition or up to 10 mg/mL for liposome preparation. Native and EAE-modified biomimetic myelin compositions were prepared according to previously reported cytoplasmic and periplasmic leaflet molar ratios (Table S1).[1]

#### Deposition of Asymmetric Supported Myelin Bilayers (SMB)

Solid supports were single crystals of silicon (8 × 5× 1 $cm^3$) polished on one large face. They were cleaned before use with chloroform, acetone, ethanol and pure water in that sequence. They were finally treated by UV–ozone (Jelight-type) for 15 min immediately prior to use.

The inner monolayer was transferred via LB method and the outer monolayer via the LS technique, as described in Rondelli et al.[2] SMB were prepared using Langmuir troughs (KSV NIMA, Biolin Scientific, Espoo, Finland) at the Partnership for Soft Condensed Matter (PSCM, Grenoble, France) and at the ISIS Neutron and Muon Source (Rutherford Appleton Laboratory,

Harwell Campus, Didcot, UK). For the LB deposition, a lipid monolayer corresponding to the periplasmic lipidic composition—comprising native myelin lipids and protiated ovine cholesterol—was spread from the organic lipid solution onto the air–water interface using ultrapure, non-buffered water as the subphase. The monolayer was then compressed to a final surface pressure of 35 mN/m, which was selected as it approaches the collapse point of myelin lipid monolayers, corresponding to the maximal packing and lateral stress conditions similar to those in in-vivo membranes.[3] A silicon wafer initially immersed (before the lipid spreading) into the subphase was vertically lifted through the monolayer at 4 mm/min under constant surface pressure. After trough cleaning and subphase renewal, the cytoplasmic leaflet (containing either protiated or deuterated cholesterol) was deposited at the air–water interface and compressed under identical conditions. Prior to LS transfer, a custom-built levelling system was used to align the substrate parallel to the air-water interface. LS transfer was performed by lowering the LB-coated crystal onto the new monolayer at 3 mm/min and finally closing the solid-liquid flow cell used for NR measurements submerged at the base of the trough.
All depositions were carried out at 15 °C under unbuffered aqueous conditions, and samples were stored under the same conditions until NR measurements, which were performed at 15–37 °C.

**Liposomes Preparation**

Lipid mixtures dissolved in organic solvent were evaporated under an argon stream, followed by vacuum annealing overnight at 50 °C. The dried lipids were rehydrated in $D_2O$ – MOPS buffer (pD = 7.4) and shaken until fully detached from the glass surface. Residual material was removed by pipetting if necessary. The mixture was sonicated for 30 minutes at 40 °C and subjected to five freeze–thaw cycles. To remove large aggregates, it was centrifuged through a 0.45 μm filter at 10,000 g for 10 minutes. The final solution was extruded twenty times through a 100 nm membrane at 50 °C.

**NR Measurements**

NR measurements were conducted at two complementary time-of-flight instruments: INTER[4] at the ISIS Neutron and Muon Source (Rutherford Appleton Laboratory, Didcot, UK), and FIGARO[5] at the Institut Laue-Langevin (ILL, Grenoble, France). On INTER, the accessible wavelength range spanned from 1.5 to 17 Å with a momentum transfer ($Q$) resolution ($\Delta Q/Q$) of 3.5%, using incident angles of 0.7° and 2.3°. On FIGARO, measurements were performed using a wavelength band of 2 to 20 Å and a resolution of 7%, with incident angles of 0.7° and

3.0°. In both setups, beam footprints were adapted to the silicon substrate dimensions and defined by slit geometry and beam divergence, yielding illuminated areas of approximately 60 mm in length and 35 mm (INTER) or 25 mm (FIGARO) in width. All measurements were carried out in time-of-flight mode, with beam alignment and data acquisition optimized to ensure consistent $Q_z$ coverage across both instruments.

Both symmetric bicytoplasmic and asymmetric SMB were prepared via sequential LB and LS depositions as described above and then sealed within the sample cell prior to beamline installation. Once in place, the cell was connected to the automated solvent exchange system. Solvent contrast variation was performed using an automated HPLC pump at a flow rate of 1 mL/min for a total exchange volume of 20 mL. Initially, bare silicon substrates were characterized in four isotopic contrasts using the same MOPS buffer composition (10 mM, pD = 7.4) prepared in different $H_2O/D_2O$ mixtures having specific neutron scattering length densities (SLD): 100% $D_2O$ (SLD: $6.35 \times 10^{-6}$ Å$^{-2}$), '4-matched water' 66% D2O/ 34% H2O (4MW; SLD: $4.00 \times 10^{-6}$ Å$^{-2}$), 'silicon-matched water' 38% D2O/ 62% H2O (SMW; SLD: 2.07 $\times 10^{-6}$ Å$^{-2}$), and 100% $H_2O$ (SLD: $-0.56 \times 10^{-6}$ Å$^{-2}$). The SMB were then characterized under the same four contrasts. MBP was incubated at $c_{MBP}$ = 0.10 mg mL$^{-1}$ for 30 min before flushing 20 mL of buffer to remove unbound protein following the protocol described in Krugmann et al. [6]. The resulting bilayer–protein complexes were measured in $D_2O$ contrast. Finally, liposomes were injected to the SMB/MBP coated wafers to form double bilayers (DBL) which were also measured in $D_2O$.

**Safety Considerations.** All procedures involving volatile organic solvents (chloroform and methanol) were carried out in a certified fume hood with appropriate personal protective equipment. UV–ozone treatment was performed following manufacturer instructions and using eye/skin protection. $D_2O$ buffers and MOPS solutions were handled using standard laboratory practices for non-hazardous reagents. No unexpected, new, or significant hazards were encountered in this work.

| Lipid type | $\phi_{\text{nat, cyt}}$ (%) | $\phi_{\text{nat, per}}$ (%) | $\phi_{\text{EAE, cyt}}$ (%) | $\phi_{\text{EAE, per}}$ (%) |
|---|---|---|---|---|
| Phosphatidylcholine (PC) | 25.9 | 18.2 | 20.1 | 13.1 |
| Phosphatidylethanolamine (PE) | 29.0 | 9.0 | 32.9 | 9.6 |
| Phosphatidylserine (PS) | 7.0 | 1.1 | 7.4 | 6.9 |
| Sphingomyelin (SM) | 6.2 | 4.2 | 2.2 | 1.3 |
| Cholesterol | 31.9 | 33.5 | 37.4 | 38.3 |
| Cerebrosides | - | 24.4 | - | 25.2 |
| Sulfatides | - | 9.6 | - | 5.6 |

**Table S1:** Native and EAE-modified cytoplasmic/periplasmic molar lipid fractions (%). Values are target compositions used to formulate monolayers/bilayers.

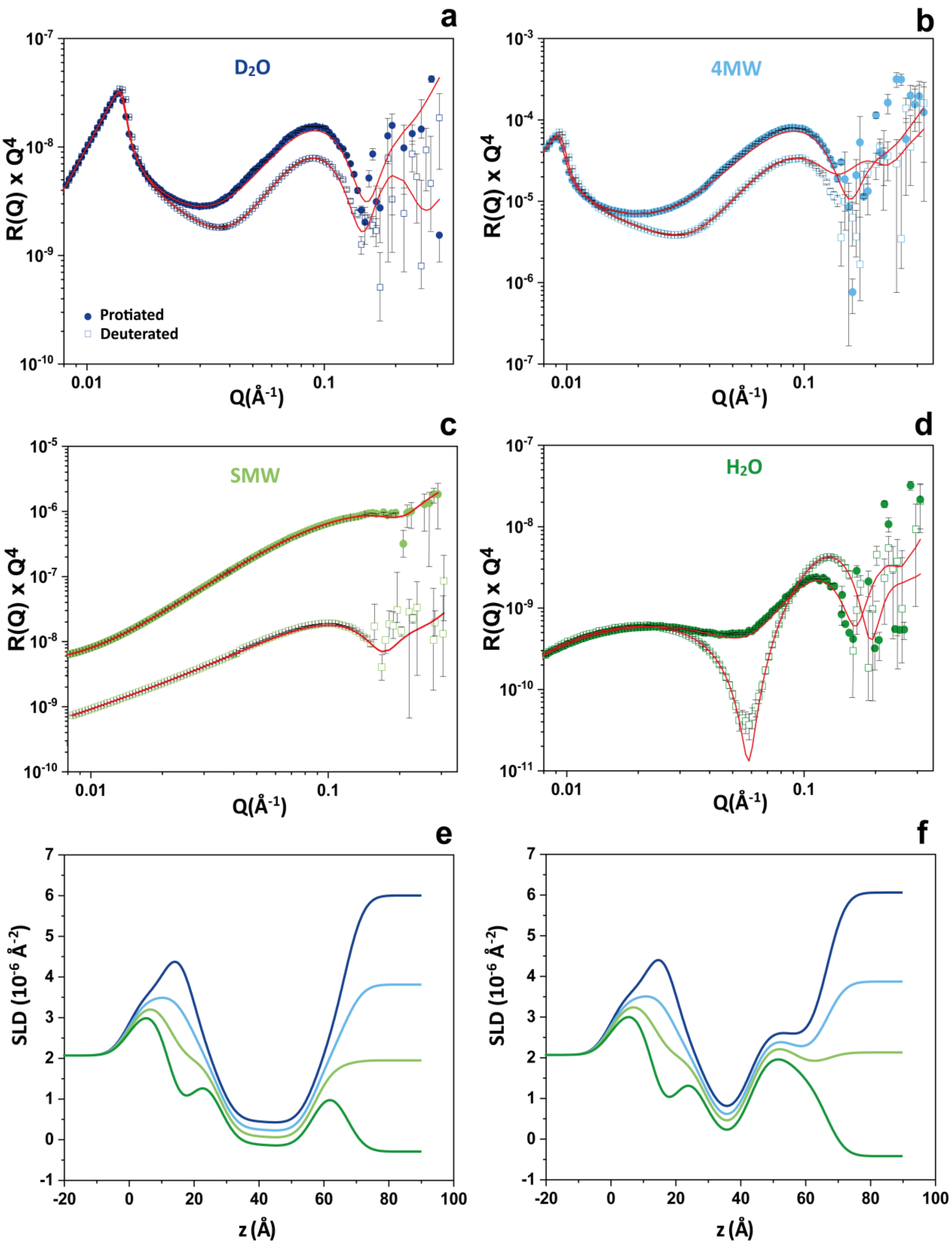


**Figure S1:** Neutron reflectivity (NR) and global co-refinement fits for an EAE-modified asymmetric SMB with either a protiated or a deuterated cytoplasmic leaflet under four solvent contrasts: (a) $D_2O$, (b) 4MW, (c) SMW, and (d) $H_2O$. Symbols denote experimental data—filled circles: protiated, open squares: deuterated—and colours encode the solvent contrast ($D_2O$: dark blue; 4MW: light blue; SMW: olive; $H_2O$: green). Red solid lines are the best co-refinement fits. Error bars represent 1σ. (e,f) Reconstructed SLD profiles (same colour coding).

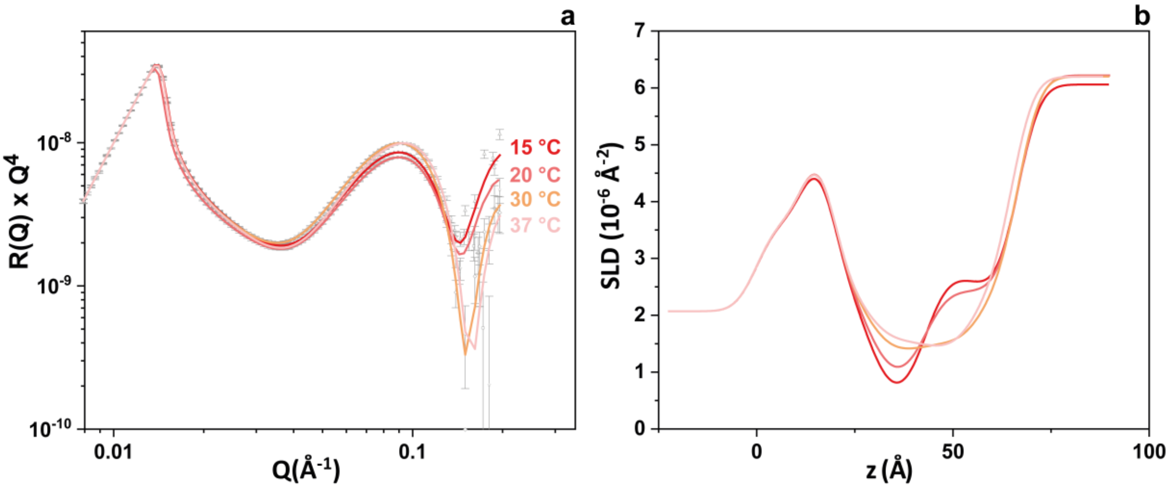


**Figure S2:** Temperature-dependent evolution of an asymmetric EAE-modified SMB with a deuterated cytoplasmic leaflet: (a) reflectivity in $D_2O$ at 15, 20, 30 and 37 °C; (b) SLD profiles.

| ***T*** (°*C*) | ***t*** (Å) | | | |
|---|---|---|---|---|
| | **Native** | | | |
| | **water** | $p_t$ | $c_t$ | **SMB** |
| **15 °C** | 4.6±1.1 | 16.2±1.1 | 16.3±1.4 | 53.1±6.5 |
| **20 °C** | 4.2±0.8 | 16.0±1.0 | 16.4±1.1 | 53.0±6.1 |
| **30 °C** | 4.9±0.7 | 15.4±1.8 | 15.2±1.0 | 51.2±6.8 |
| **37 °C** | 5.3±1.5 | 14.6±1.1 | 14.0±1.9 | 49.2±7.0 |
| | **EAE-modified** | | | |
| | **water** | $p_t$ | $c_t$ | **SMB** |
| **15 °C** | 4.4±0.8 | 14.1±1.7 | 13.8±1.3 | 48.2±5.7 |
| **20 °C** | 4.6±1.2 | 14.2±1.3 | 14.0±1.4 | 48.5±5.0 |
| **30 °C** | 3.9±0.8 | 13.9±1.2 | 14.1±1.5 | 48.3±5.0 |
| **37 °C** | 5.2±1.6 | 13.3±1.4 | 13.1±1.1 | 46.7±5.2 |

**Table S2:** Temperature series of thickness ($t$) parameters. Interfacial-water thickness, leaflet acyl-chain thicknesses ($p_t$, $c_t$), and total bilayer thickness (SMB = $p_h + p_t + c_t + c_h$) for native and EAE-modified asymmetric SMB with a deuterated cytoplasmic leaflet at 15, 20, 30 and 37 °C. Uncertainties are 1σ from the fit covariance; totals use full covariance propagation.

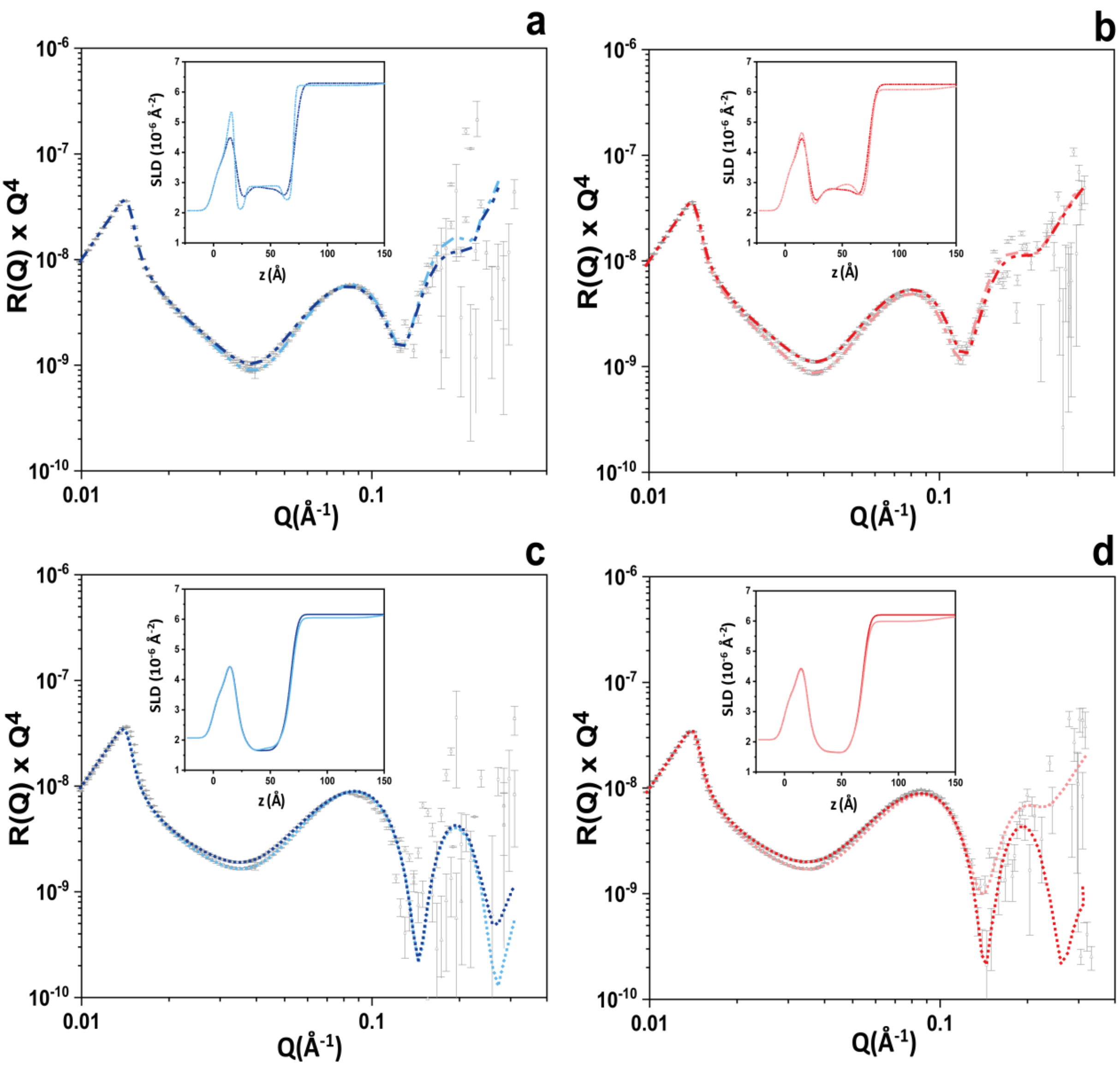


**Figure S3:** NR data and co-refinement fits for bicytoplasmic and cholesterol-redistributed SMB. Panels (a,b): bicytoplasmic SMB at 15 °C for native (a) and EAE-modified (b) compositions; fitted curves are shown as dash–dot lines. Panels (c,d): cholesterol-redistributed SMB at 37 °C for native (c) and EAE-modified (d); fits are shown as dotted lines. Experimental data are plotted as grey symbols—squares for the bare membranes and triangles after MBP addition. In each panel, dark and light shades correspond respectively to the membrane before and after protein binding; insets display the corresponding SLD profiles with matching colour coding. Error bars represent 1σ uncertainties.

| | Native | | EAE-modified | |
|---|---|---|---|---|
| | **Bicytoplasmic** | **37 °C** | **Bicytoplasmic** | **37 °C** |
| $t_{SMB}$ (Å) | 53.0 ± 4.8 | 49.2 ± 7.0 | 50.7 ± 5.6 | 46.7 ± 5.2 |
| $t_{SMB}$ after MBP-adsorption (Å) | 54.2 ± 5.5 | 51.3 ± 5.2 | 51.8 ± 6.6 | 48.8 ± 4.9 |
| $t_{MBP}$ (Å) | 71.9 ± 7.9 | 68.3 ± 5.2 | 68.1 ± 11.9 | 65.0 ± 9.8 |
| $c_{MBP/SMB}$ (%) | 3 ± 4 | 2 ± 5 | 5 ± 4 | 6 ± 3 |

**Table S3:** Fitted structural parameters for bicytoplasmic and cholesterol-redistributed SMB (native and EAE-modified), with uncertainties ± 1σ.
